# Estimated incidence of noma “A biologic indicator of poverty” in north central Nigeria: A retrospective cross-sectional study

**DOI:** 10.1101/549956

**Authors:** Seidu A. Bello, John A. Adeoye, Ifeoluwa Oketade, Oladimeji A. Akadiri

**Author notes:** Corresponding author: (SB). These authors contributed equally to this work (Seidu A. Bello, John A. Adeoye).

## Abstract

**Background:** Noma is a spreading and devouring disease which is believed to be native to Sub-Saharan Africa over the last decade due to poverty. Within this noma belt, most epidemiological reports regarding the disease have emanated from the north western region of Nigeria. However, our indigenous surgical mission encountered a substantial number of cases noteworthy of epidemiological representation in north central Nigeria.

**Methods:** All facial cleft and noma cases encountered within the 8 year study period were included into this study. Estimated incidence of the noma in the zone was calculated using the existing statistical model of Fieger *et al* (2003), which takes into account the expected differences based on age and location of the two patient groups using the multinomial logistic regression analysis. Period prevalence of noma was also calculated by simple division considering the population at risk of the disease in the zone.

**Findings:** A total of 770 subjects were included in this study (orofacial cleft – 692, noma – 78). The incidence estimate of noma in the north central zone was 3.2 per 1000 with a range of 2.6 – 3.7 per 1000. The period prevalence of noma was1:125,000 children. The median age of noma patients was comparatively higher than the median age of facial cleft patients. The mean age of onset of noma was 5.9 ± 8.08 years which was lower than the average age of individuals in the noma group - 29.6 ± 18.84 years.

**Conclusion:** Although noma may be more prevalent in the north western region of Nigeria; substantial number of cases is still being encountered in the north central zone which calls for urgent attention of relevant health stakeholders regarding the management and rehabilitation of individuals affected.

**AUTHOR SUMMARY:** Noma, a devouring facial disease, is commonly associated with poverty and impoverished regions of the world especially Sub-Saharan Africa which is being termed the noma belt region of the world. Although literature established that noma is indeed a neglected disease, the degree of this neglect in north central Nigeria compared to other sub-regions is in fact alarming, as no report on the disease burden have been published till date. In this light, a retrospective, cross-sectional was conducted to provide epidemiological representation to the cases encountered within an eight year period at the Cleft and Facial Deformity Foundation (CFDF), an indigenous surgical mission. The incidence of noma was estimated from the known incidence of orofacial cleft using an existing multinomial logistic regression model while the period prevalence was calculated considering the population living below poverty line in the sub-region. This study extrapolates an incidence of 3.2 per 1000 and a period prevalence of 0.05 per 1000 persons. Notable is the finding that most individuals with noma were above thirty years of age and suffered varying degree of facial disfigurement resulting from the acute phase of the disease which started in their childhood. Therefore, we advocate public awareness on the disease presentation, risk factors and sequelae in the sub-region and identify the need to bolster the efforts of existing health facilities and indigenous surgical missions in the management and rehabilitation of individuals affected.

## INTRODUCTION

Noma, alternatively known as Stomatitis gangrenosa and Cancrum oris, is derived from the Greek word “nome”, meaning “a pastureland or grazing” which is a figurative term describing the devouring nature of the disease and spread of its lesions.^[1,2]^ It occurs as a continuously spreading ulcer of the mouth and was first described clinically by Carel Baten in 1595.^[3]^ Although recently, the occurrence of noma has been linked with various immune suppressing conditions such as HIV infection, leukaemia, Non-Hodgkin’s lymphoma and cyclic neutropenia;^[4]^ it is still almost exclusively associated with extreme poverty especially in Sub-Saharan Africa.^[5,6]^

Noma has a noteworthy history, with poverty and its sequelae as the central factor. In Northwest Europe, noma became prevalent around the 18th century and was related to poverty, malnutrition, and preceding diseases most especially measles.^[6]^ At the end of that century, noma gradually disappeared in the Western world because of economic progress, which gave the poorest in the society the opportunity to feed their children sufficiently.^[6,7]^ Only during the World War II([WW II], 1939-1945) did noma reappear in concentration camps, and in the Netherlands, where the population suffered from famine towards the end of the war.^[4]^ Consequently, with noma being a thing of the past in developed countries, the twentieth century witnessed increased discovery and alarming reports of a huge burden of the disease in Africa as documented from the experience of Europeans on volunteer surgical missions to the continent.^[8,9]^ More recently, most noma patients can probably be found in the impoverished savannah region directly south of the Sahara desert in Africa, which is known as the world’s “noma belt” extending from Senegal to Ethiopia.^[6]^

With comparatively little attention to the scourge of Noma in Africa by the world bodies, one major problem identified is the lack of local epidemiological data on the disease in affected regions.^[7]^ A prevalence of 770,000 persons, who survived the disease with devastating sequelae was first reported in 1997, followed in 1998 with the World Health Report by Bourgeois & Leclercq who found an estimated incidence of 140,000 cases.^[10,11]^ In Nigeria, Barmes et al^[12]^ estimated a case incidence of 1:1250, alongside worse case incidence reports in Niger and Senegal. Furthermore, Fieger et al^[7]^ in an estimation of the incidence of Noma in Sokoto, North West Nigeria, reported an incidence of 6.4 per 1000 children. Extrapolation of this incidence to the developing countries bordering the Sahara Desert (the noma belt of the world) gives an incidence of 25,600 for that region and a global incidence of 30,000–40,000.^[7]^

Reports by Marck et al^[13]^ revealed that surgical rehabilitation of noma survivors in Nigeria started in 1996 in Sokoto State, north west region of the county. In fact, the study by Fieger et al in 2003 was based on the observations of the surgical outreach between 1996 and 2001 in the same region.^[7]^ A similar free surgical outreach programme was commenced in 2010 by an indigenous surgical mission – CLEFT & FACIAL DEFORMITY FOUNDATION (CFDF) with an aim of awareness creation and surgical management of several craniofacial and dental conditions in a region challenged with the dearth of secondary and tertiary health institutions. Epidemiological data is important for planning and prioritisation of service delivery and hence continuous reappraisal is always very important. Although it is widely believed that the noma scourge is exclusive to northwest Nigeria as evidenced by the number of reports that have emanated from the region and the establishment of an health institution – Noma Children Hospital, solely concerned with treatment of acute stages of the disease and rehabilitation of survivors. Cases of noma are also being encountered and successfully managed by our surgical mission within north central Nigeria in recent times. The aim of the present study therefore is to estimate the incidence of noma in north central, Nigeria using the same statistical methods utilized by Feiger et al^[7]^ with an additional emphasis on associated factors and patients’ demographics in comparison to the orofacial cleft presentations in the same sub-region.

## MATERIALS AND METHODS

This was a retrospective cross-sectional, epidemiological study of facial clefts and noma patients seen at various free surgical outreach programme locations in north central Nigeria between 2010 and 2018. Cleft and Facial Deformity Foundation (CFDF) is an indigenous surgical mission that has its focus on free surgical care for individuals with orofacial diseases since 2010. The organization is based in the north central geo-political zone of the country where it has been carrying out outreach programmes in different locations of the zone.

Comprehensive information were obtained from stored records of patients encountered in all surgical outreach programmes organized in north central Nigeria from June 2010 till September 2018. All cases diagnosed as facial clefts and nomas were included in this study, and this comprised adults and children alike while other lesions of head and neck region were excluded. The information obtained from the records included the bio-data, definitive diagnoses and description of the lesions at presentation. Distance between patients’ location of residence and the health institutions where the surgical outreaches were conducted were also estimated manually for both patient groups (facial cleft and noma). Other information obtained solely from the noma group included the age of disease onset, proximity of residence to livestock (cattle, pigs, horses etc), number of siblings as well as information ascertaining whether the patient was residing with an extended family member around the time of onset of the disease. Written permission was obtained from the patients for the use of their pictures for educational and research purposes. Resultant from the prior experience of members of the Cleft and Facial Deformity Foundation Data Management Team (CFDF-DMT), most variables perceived to be missing from a number of patient records on collection (due to several factors) were obtained from recall visits usually organized about 3-months following the surgical outreach program. At the time of information collation, other participants with missing data whose information could not be obtained from the then information provider were further excluded from the study. This was necessary to reduce output uncertainty in the study. However, only information regarding the age of noma onset were left unanswered if the individuals or their informants could not approximately estimate the time.

Data obtained from the study was analyzed using Statistical Package for Social Sciences (SPSS) version 23.0 (IBM Corp Armonk, NY, USA). Descriptive statistics such as frequencies, mean and standard deviation were explored as required for quantitative and categorical variables. The normality of the distribution was ascertained using the Shapiro-Wilk’s test. Difference between quantitative variables was determined using the Mann-Whitney U test. Relationships between categorical variables were determined using the Pearson’s Chi-square test. The prevalence of the disease was calculated with the number of noma cases seen within the study period as numerator and the population at risk as denominator multiplied by 1000. The population at risk included only 45.7% of individuals residing below poverty line in the north central zone of Nigeria.^[14]^

Incidence of noma was calculated with the multinomial logistic regression analysis model of Feiger et al^[7]^ according to the age of the individuals and their distance between their residence and surgical outreach centre, so as to take into account the predicted odds between the two patient groups. With the absence of recent incidence values for facial cleft, we utilized the reference data of Iregbulem et al^[15]^ (0.4 per 1000) and a noma mortality rate of 90%^[5]^; which were also used in the study by Feiger et al.^[7]^ The statistical values of all statistical tests used were set to 5% (p<0.05). Ethical approval for the study was obtained from the Research Ethics review board of the International Craniofacial Academy and written consents were obtained from participants themselves or their legal guardians (for paediatric participants) as at the time of data collection.

## RESULTS

Within the eight year study period, our indigenous surgical mission encountered 708 cases of orofacial cleft (OFC) and 78 noma cases at various centres across north central Nigeria. Sixteen (16) OFC participants were excluded due to one or more missing required data in their records; finally, 692 individuals were included for the analyses.

Age and sex variables of both groups of patients were not normally distributed (Shapiro-Wilk’s test, p<0.05). The average age of individuals with orofacial cleft and noma were 8.5 ± 10.94 years and 29.6±18.84 years respectively. More OFC patients were less than 5 years of age (n = 376, 54.3%) while majority of noma patients (n = 43.6, 34.3%) were above the age of thirty; the difference was statistically significant (p = 0.001) (Table 1). Figure 1 shows the box-plot of the ages of orofacial cleft and noma patients in this study. Each element of the box and whisker plot in terms of their ages was observed to be higher in noma patients than in individuals with OFC.

**Table 1:** Age and sex Characteristics of orofacial cleft and noma patients

|  | Cleft<br>N (%) | Noma<br>N (%) | P Value |
| --- | --- | --- | --- |
| Age (in years) |  |  |  |
| <5 | 376 (54.3) | 6 (7.7) |  |
| 5 – 10 | 118(17.1) | 7 (9.0) |  |
| 11 – 15 | 58 (8.4) | 11 (14.1) | U = 7553.500; |
| 16 – 20 | 49 (7.1) | 7 (9.0) | P = 0.001* |
| 21 – 25 | 26 (3.8) | 7 (9.0) |  |
| 26 – 30 | 35 (5.1) | 6 (7.7) |  |
| >30 | 30 (4.3) | 34(43.6) |  |
| Sex |  |  |  |
| Male | 343 (49.6) | 42 (53.8) | $\chi^2 = 0.514$; df = 1;<br>0.474 |
| Female | 349 (50.4) | 36 (46.2) |  |

\*Statistical significant difference; $P < 0.05$ ; Mann-Whitney U test

**Figure 1:**
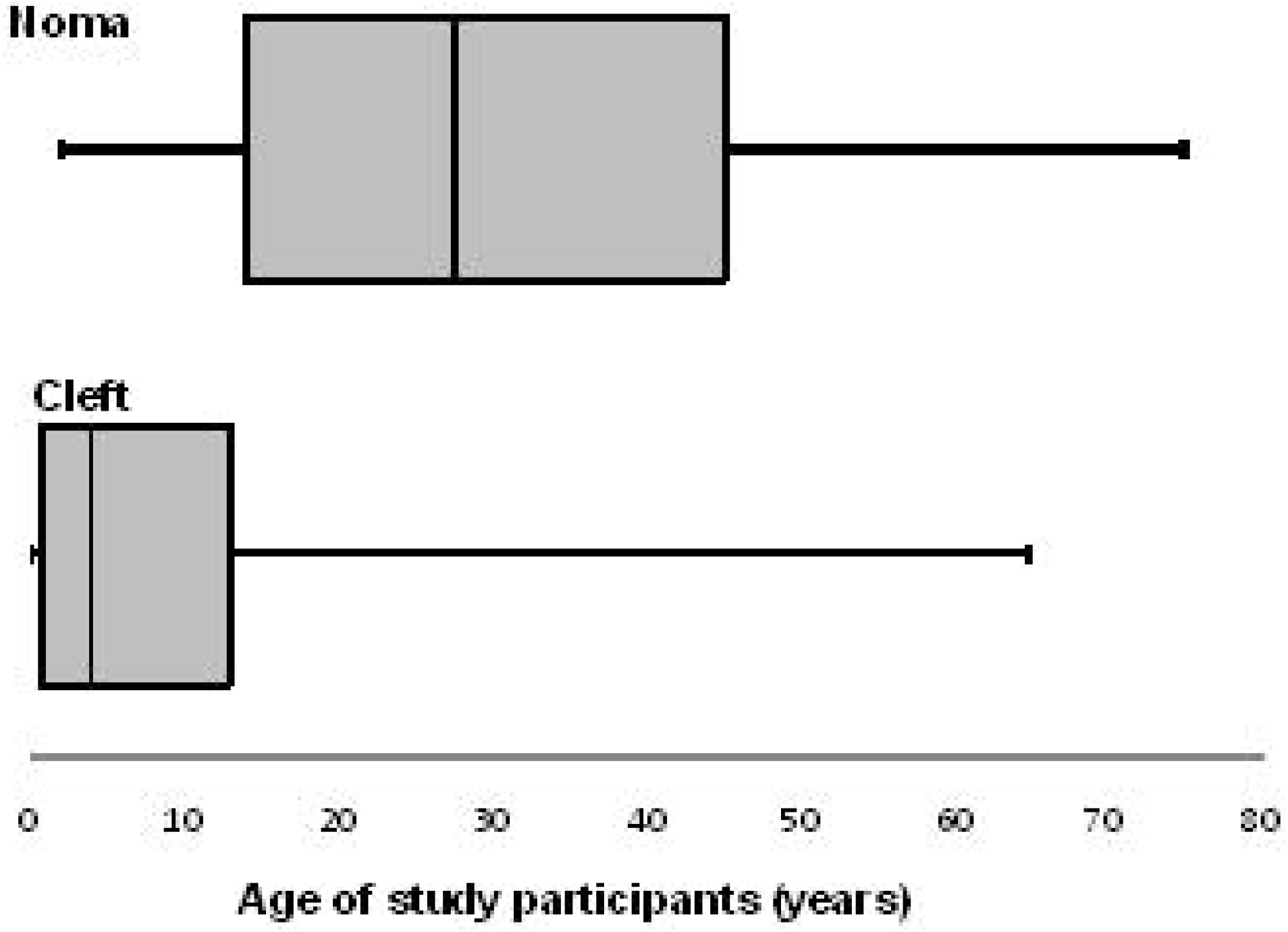
Picture showing a diagnosed case of noma at one of the surgical outreach programmes.

Figure 1: Box and Whisker Plot of the ages of orofacial cleft and noma patients

The mean distance between patients’ residence and location of the health facility used for the surgical outreach was 126.9±127.79 km for OFC patients and 124.8±96.713 km for noma patients. In addition, most participants in both groups (n = 284, 41.0% in OFC group and n = 41, 52.6% in noma group) mostly resided in locations approximately 30 – 100km from the health facility used (Table 2).

**Table 2:**
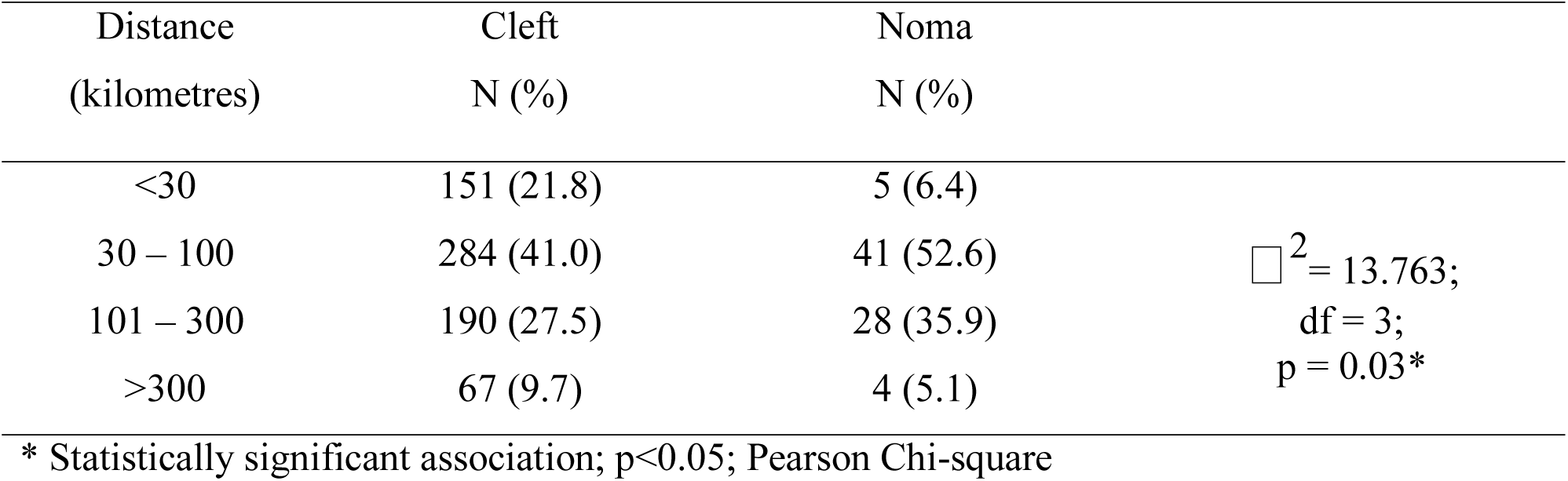
Distribution of the distance between patients’ residence and health facility in both groups

| Distance<br>(kilometres) | Cleft<br>N (%) | Noma<br>N (%) |  |
| --- | --- | --- | --- |
| <30 | 151 (21.8) | 5 (6.4) |  |
| 30 – 100 | 284 (41.0) | 41 (52.6) | $\chi^2 = 13.763$;<br>df = 3;<br>$p = 0.03^*$ |
| 101 – 300 | 190 (27.5) | 28 (35.9) |  |
| >300 | 67 (9.7) | 4 (5.1) |  |

\* Statistically significant association; $p < 0.05$ ; Pearson Chi-square

The age of onset of noma in this study ranged from 3 months to 30 years of age with an average of 5.9± 8.08 years. Most patients or informants (n = 57, 73.0%) could not ascertain the age of onset of the disease; however, 9.0% of the patients (n = 12), which accounted for most of the patients that could fairly estimate the ages of onset of noma, claimed the symptoms started when they were about two years old (Table 3).

**Table 3:** Distribution of the age of onset of noma

| Age (years) | Frequency (%) |
| --- | --- |
| 0 – 1 | 5 (6.4) |
| 1 – 2 | 7 (9.0) |
| 3 – 7 | 5 (6.4) |
| 14 | 1 (1.3) |
| 20 – 30 | 3 (3.9) |
| Missing/NA | 57 (73.0) |

The period prevalence of OFC and noma calculated in this study were 0.05 per 1000 and 0.008 per 1000 respectively. For the noma group, calculated prevalence for males and females were 0.009 per 1000 and 0.007 per 1000. Table 4 shows that the estimated incidence calculated using the relevant age group (10 – 30 years) and relevant distance to the health facilities (>100 km) was 3.2 per 1000, with a range of 2.6 – 3.7 per 1000. Pertinent information collected regarding other risk factors associated with noma included the number of siblings in the family, being raised by extended family members (especially grandparents) and the proximity of household residence to livestock. The number of siblings of individuals in the noma group ranged from 3 to 18 in total with an average of 8.6 ± 5.06. Furthermore, 85.7% (n = 67) answered positively that they lived in close proximity to livestock or even reared them while only 18.8% (n = 15) admitted to residing with extended relatives at the time of onset of noma disease.

**Table 4:** Estimated incidence of noma according to relevant ages and location of patients

| Age (years) | Distance of patients' residence to health facility (km) | 95% CI | p-value | Estimated incidence | Weighted average of estimated incidence | Cumulative average of estimated incidence |
| --- | --- | --- | --- | --- | --- | --- |
| 10 – 15 |  | 1.73, 13.95 | 0.001 |  |  | 3.2 |
|  | 100 – 300 | 0.12, 1.0 | 0.05 | 3.9 | 3.6 |  |
|  | >300 | 0.13, 1.13 | 0.08 | 3.1 |  |  |
| 16 – 20 |  | 1.32, 12.44 | 0.001 |  |  |  |
|  | 100 – 300 |  |  | 2.8 | 2.6 |  |
|  | >300 |  |  | 2.3 |  |  |
| 21 – 30 |  | 1.60, 14.10 | 0.001 |  |  |  |
|  | 100 – 300 |  |  | 3.5 | 3.3 |  |
|  | >300 |  |  | 2.7 |  |  |

## DISCUSSION

Our study shifts major attention from the norm of conducting noma epidemiological surveys in the north western region of Nigeria in the last decade to the north central region which consists of seven member states (Plateau, Kwara, Nasarawa, Niger, Benue, Kogi and the Federal Capital Territory). Over the study period, we have discovered that there has been increasing number of cases noteworthy of epidemiologic representation so as to adequately characterize the disease burden in this region which has the third-highest number of citizens living below the poverty line in Nigeria.^[14]^

The combined estimated incidence of noma in north central Nigeria from 2010 – 2018 is 3.2 per 1000 although ranging from 2.6 to 3.7 per 1000 depending on the age category. This is approximately half less than the calculated incidence of noma reported by Feiger et al^[7]^ from 378 noma patients encountered between 1996 – 2001. In the later study, the estimated incidence was 6.4 per 1000 with a range of 4.4 – 8.5 per 1000 in individuals between 10 – 30 years of age. Our lower incidence estimate may be clearly attributed to the wide variation in poverty indices of the north western and north central region of the country over several years, with the north western region serially recording the highest number of individuals living below poverty line in the country (>80.0%).^[14]^ By inference therefore, the north central region may have less residents with severe malnutrition, unsafe drinking water, poorer sanitation practices and limited access to proper healthcare when compared to the north western sub-region. Our calculated incidence was also lower than what was obtained by Denloye et al^[16]^ in south west Nigeria from 1986 – 2000 with an incidence rate of 7.0 per 1000 cases in individuals between 1 – 12 years of age. In further comparison of our findings with earlier reports from other regions in the noma belt of the world, we observed that our estimated incidence was higher than the case incidence reported from Niger Republic (1.34 per 1000) and Senegal (0.7 – 1.2 per 1000) by Barmes et al^[12]^ among children aged 0 – 6 years. However, this comparison may be flawed since at the time of data extrapolation, a seemly unrealistic mortality rate of 70% was utilized for the incidence estimation in their study; this observation was also corroborated by the reports of Fieger et al^[7]^.

The period prevalence of noma and orofacial cleft in this study was 0.008 per 1000 and 0.05 per 1000 respectively. Regarding the socio-demographic characteristics of study groups, the mean age of individuals with noma was 29.6 ± 18.84 years which was significantly higher than the average age of OFC patients with most individuals in the facial cleft group being less than 5 years of age. This finding is consistent with the age distribution of both facial cleft and noma patients reported by Fieger et al^[7]^ in north western Nigeria. However, our calculated average age of noma patients was markedly higher than the reports of Denloye et al^[15]^ in south-western Nigeria and Enwonwu et al^5^ across various Nigerian sub-regions. In the later study involving over 1000 Nigerian children from different geo-political zones, a mean age of 5.96 ± 2.62 years was obtained, with most of the study subjects falling below the age of 13. Although the average age of noma patients in this study was above 25 years, the mean age of noma onset was 5.9 ± 8.08 years which implied that most of the patients encountered were not in the acute phase of the disease and perhaps suffered varying degrees of cosmetic disfigurement which led to their presentation at the surgical outreaches. Pertaining to the disease sequelae, this may also mean that the acute stage of the disease occurs commonly in children, which is consistent with the report of Braimah et al^[17]^ in north western Nigeria. It is noteworthy to state that two-thirds of noma patients could not provide an approximate age of onset of the disease which may indicate that the natural disease process in these patients were perceived as unremarkable or that patients were too young at the time of onset to remember. With majority of noma patients presenting at >30 years while available data on disease onset suggests greater frequency in early childhood, it can be inferred that most survivors of the acute phase lived without proper surgical rehabilitation. Compared to noma, the proportion of OFC in adult patients is significantly low. This can be adduced to the increased awareness and massive foreign investment in providing free surgical correction of congenital orofacial cleft deformities in Nigeria and training of manpower for this purpose mainly spearheaded by the SmileTrain organization (New York, USA) in the last two decades; this is yet to happen with noma. In addition to age, we observed that the sex of both noma and OFC groups were fairly distributed and play little or no role in describing the incidence of both disease entities.

Other than poverty, some other risk factors for the disease were also investigated. Our study observed that close proximity of noma patients to livestock around the time of disease onset may be a risk factor for the disease. This observation is similar to the reports of Enwonwu et al^3^, Barmes et al^[12]^ and Baratti-Meyer et al^[18]^, and may strengthen the theory that domestic livestock breeding creates an unhygienic environment and dwelling in proximity to such may play a major role in the disease process. However, further studies are still required to confirm this observation. It was also observed that 18.8% of noma patients were being catered for by extended relatives at the time of disease onset. Farley et al^[19]^ in a case-control study involving 74 cases and 222 controls in north west Nigeria, associated “caretaker” (i.e third party carer) as a factor that may influence the risk of developing noma. This was supported by Adeola et al^[20]^ as reported in a cases series involving five subjects managed for acute phase of noma in north west Nigeria. In that study, they asserted there was increasing number of noma cases in the region associated with lack of direct maternal care after children were weaned. This observation was not common in our study where only 18.8% of noma patients were being catered for by extended relatives at the time of disease onset. Although this proportion may be considerable, it was not statistically significant to corroborate the observation of the earlier authors. Hence, it may require further studies to determine the plausibility of this assertion. Another important observation is the travel distance to access health care for patients in the north central sub-region. In this study, patients had to travel above 100km on the average to access care from our surgical team this is partly because there were no hospitals with required facilities and competence in closer proximity and partly because patient could not afford the cost of treatment and additional cost of transportation. This burden was relieved by the gesture of our surgical mission.

Considering that our study sought out to primarily determine the estimated incidence and period prevalence in north central Nigeria, these results that have emanated are specific to characterizing the burden of noma in this sub-region. However, the statistical methods for the calculations done were adapted from an earlier study and could as well be applied to other regions of the country or in other locations. Another limitation encountered is the use of non-probability sampling methods to arrive at the sample size employed for the analysis of other variables such as age of noma onset and associated risk factors for the disease which may have introduced selection bias to the study.

## CONCLUSION

The prevalence and estimated incidence of noma in north central Nigeria is 1 in 125,000 persons and 3.2 per 1000 respectively while the prevalence of OFC was 1 in 20,000. Although the noma scourge is deemed prevalent in north west Nigeria and Sokoto state in particular, substantial number of cases is being encountered in the north central zone due to disease onset in native individuals. Hence, efforts should be intensified in terms of public awareness regarding the disease in the north central sub-region as well as education of relevant health care practitioners in the management of the condition. With majority living with noma or post-noma defect at >30 years of age in this sub-region, it is clear that attention to surgical rehabilitation in the region is suboptimal. On the other hand, the proportion of adult patients living with congenital orofacial cleft is dwindling due to the tremendous attention OFC has got in the last two decades. It is therefore imperative that in the absence of any health facility solely dedicated to the management and rehabilitation of noma patients in the region (unlike the northwest), existing secondary health centres and non-governmental indigenous surgical missions in the zone should be better equipped to mitigate the increasing disease burden especially as the poverty index of the zone and country is increasing.

## Supporting information

STROBE Checklist

## ABBREVIATIONS

WWII –: World War II
pCFDF –: Cleft and Facial Deformity Foundation
SPSS –: Statistical package for the Social Sciences
OFC –: Orofacial Cleft

## SUPPORTING INFORMATION LEGENDS

S1 Checklist: STROBE Checklist

