## Supplementary material for "Estimated incidence of noma “A biologic indicator of poverty” in north central Nigeria: A retrospective cross-sectional study": STROBE Checklist

|  | Item No | Recommendation | Manuscript Section | Paragraph number |
| --- | --- | --- | --- | --- |
| **Title and abstract** | 1 | (*a*) Indicate the study’s design with a commonly used term in the title or the abstract | Cover page | Paragraph 1 |
|  |  | (*b*) Provide in the abstract an informative and balanced summary of what was done and what was found | Abstract | Sub-headings 1-4 |
| Introduction | | |  |  |
| Background/rationale | 2 | Explain the scientific background and rationale for the investigation being reported | Introduction | Paragraphs 1-4 |
| Objectives | 3 | State specific objectives, including any prespecified hypotheses | Introduction | Paragraph 4 |
| Methods | | |  |  |
| Study design | 4 | Present key elements of study design early in the paper | Materials and Methods | Paragraph 1 |
| Setting | 5 | Describe the setting, locations, and relevant dates, including periods of recruitment, exposure, follow-up, and data collection | Materials and Methods | Paragraphs 1-2 |
| Participants | 6 | (*a*) Give the eligibility criteria, and the sources and methods of selection of participants | Materials and Methods | Paragraph 2 |
| Variables | 7 | Clearly define all outcomes, exposures, predictors, potential confounders, and effect modifiers. Give diagnostic criteria, if applicable | Materials and Methods | Paragraph 2 |
| Data sources/ measurement | 8* | For each variable of interest, give sources of data and details of methods of assessment (measurement). Describe comparability of assessment methods if there is more than one group | Materials and Methods | Paragraph 2 |
| Bias | 9 | Describe any efforts to address potential sources of bias | Materials and Methods | Paragraph 2 |
| Study size | 10 | Explain how the study size was arrived at | Materials and Methods | Paragraph 2 |
| Quantitative variables | 11 | Explain how quantitative variables were handled in the analyses. If applicable, describe which groupings were chosen and why | Materials and Methods | Paragraph 3 |
| Statistical methods | 12 | (*a*) Describe all statistical methods, including those used to control for confounding | Materials and Methods | Paragraphs 3-4 |
|  |  | (*b*) Describe any methods used to examine subgroups and interactions | Materials and Methods | Paragraphs 3-4 |
|  |  | (*c*) Explain how missing data were addressed | Materials and Methods | Paragraph 2 |
|  |  | (*d*) If applicable, describe analytical methods taking account of sampling strategy | Not applicable |  |
|  |  | (*e*) Describe any sensitivity analyses | Materials and Methods | Paragraph 2 |
| Results | | |  |  |
| Participants | 13* | (a) Report numbers of individuals at each stage of study—eg numbers potentially eligible, examined for eligibility, confirmed eligible, included in the study, completing follow-up, and analysed | Results | Paragraph 1 |
|  |  | (b) Give reasons for non-participation at each stage | Results | Paragraph 1 |
|  |  | (c) Consider use of a flow diagram | Not considered |  |
| Descriptive data | 14* | (a) Give characteristics of study participants (eg demographic, clinical, social) and information on exposures and potential confounders | Results | Paragraph 1  Table 1 |
|  |  | (b) Indicate number of participants with missing data for each variable of interest | Results | Paragraph 1,4 |
| Outcome data | 15* | Report numbers of outcome events or summary measures | Results | Paragraph 5  Table 4 |
| Main results | 16 | (*a*) Give unadjusted estimates and, if applicable, confounder-adjusted estimates and their precision (eg, 95% confidence interval). Make clear which confounders were adjusted for and why they were included | Results | Table 4 |
|  |  | (*b*) Report category boundaries when continuous variables were categorized | Results | Tables 1,3-4 |
|  |  | (*c*) If relevant, consider translating estimates of relative risk into absolute risk for a meaningful time period | Not relevant | Not relevant |
| Other analyses | 17 | Report other analyses done—eg analyses of subgroups and interactions, and sensitivity analyses | Results | Paragraph 2 |
| Discussion | | |  |  |
| Key results | 18 | Summarise key results with reference to study objectives | Discussion | Paragraph 2 |
| Limitations | 19 | Discuss limitations of the study, taking into account sources of potential bias or imprecision. Discuss both direction and magnitude of any potential bias | Discussion | Paragraph 5 |
| Interpretation | 20 | Give a cautious overall interpretation of results considering objectives, limitations, multiplicity of analyses, results from similar studies, and other relevant evidence | Discussion | Paragraphs 2-5 |
| Generalisability | 21 | Discuss the generalisability (external validity) of the study results | Discussion | Paragraph 5 |
| Other information | | |  |  |
| Funding | 22 | Give the source of funding and the role of the funders for the present study and, if applicable, for the original study on which the present article is based | Funding Statement | Paragraph 1 |

*Give information separately for exposed and unexposed groups.

**Note:** An Explanation and Elaboration article discusses each checklist item and gives methodological background and published examples of transparent reporting. The STROBE checklist is best used in conjunction with this article (freely available on the Web sites of PLoS Medicine at http://www.plosmedicine.org/, Annals of Internal Medicine at http://www.annals.org/, and Epidemiology at http://www.epidem.com/). Information on the STROBE Initiative is available at www.strobe-statement.org.
